# LINKS: Scaffolding genome assemblies with kilobase-long nanopore reads

**DOI:** 10.1101/016519

**Authors:** René L. Warren, Benjamin P. Vandervalk, Steven J.M. Jones, Inanç Birol

## Abstract

**Motivation:** Owing to the complexity of the assembly problem, we do not yet have complete genome sequences. The difficulty in assembling reads into finished genomes is exacerbated by sequence repeats and the inability of short reads to capture sufficient genomic information to resolve those problematic regions. Established and emerging long read technologies show great promise in this regard, but their current associated higher error rates typically require computational base correction and/or additional bioinformatics preprocessing before they could be of value. We present LINKS, the Long Interval Nucleotide K-mer Scaffolder algorithm, a solution that makes use of the information in error-rich long reads, without the need for read alignment or base correction. We show how the contiguity of an ABySS *E. coli* K-12 genome assembly could be increased over five-fold by the use of beta-released Oxford Nanopore Ltd. (ONT) long reads and how LINKS leverages long-range information in *S. cerevisiae* W303 ONT reads to yield an assembly with less than half the errors of competing applications. Re-scaffolding the colossal white spruce assembly draft (PG29, 20 Gbp) and how LINKS scales to larger genomes is also presented. We expect LINKS to have broad utility in harnessing the potential of long reads in connecting high-quality sequences of small and large genome assembly drafts.

**Availability:** http://www.bcgsc.ca/bioinformatics/software/links

## 1 INTRODUCTION

Long read technology has rapidly matured over the past few years, and the benefit of long reads for genome assembly is indisputable (reviewed by Koren and Phillippy, 2014). Recently, groups have shown that *de novo* assembly of error-rich long reads into complete bacterial genomes is possible, even without prior base correction (Berlin *et al.*, 2014)

Portable long read sequencing technology is at our doorstep, thanks to leaps in microfluidics, electronics and nanopore technologies (Clarke *et al.*, 2009). Expected to be a strong contender in the kilobase-long read arena, Oxford Nanopore Technologies Ltd (ONT, Oxford, UK) promises a miniature molecule “sensor” that is currently in limited early access beta-testing phase through the MinION™ Access Programme (MAP). At the moment, sequence reads generated by the instrument have limited utility for *de novo* assembly of genomes, which is mostly due to their associated high base errors and indel rates (Quick *et al.*, 2014). Very recently, Quick *et al.* (2014) publicly released ONT *E. coli* long reads as part of the MAP. Whereas their assessment identified some of the shortcomings of the current technology, it also highlighted its great potential, including a low-cost throughput and kilobase long reads.

High quality R7 chemistry data (termed Full 2D) in the released dataset comprises reads derived from template and complementary strands. To our assessment, these reads have an average sequence identity of 77.1 +/− 10.6% (11 Mbp in 1714 reads with sequence identity of 50% or more to *E. coli* K-12 MG1655). Despite this low overall quality, there are still frequent continuous stretches of correct *k* bases in the reads when compared to the finished genome. These stretches are long enough to confer specificity, but short enough to be error free (Fig. S1, *k*=15). We have exploited this property of the sub-5X data (Fig. S2) to develop a genome scaffolding algorithm, LINKS, which extracts paired *k-mers* from the ONT reads, and uses them to link contig pairs. One advantage of the proposed implementation is its ability to iteratively refine assemblies by exploring large numbers of *k-mer* pair combinations for linking contigs.

## 2 METHODS

### 2.1 Sequence data, assembly, and scaffolding

*E. coli* K-12 substr. MG1655 Illumina MiSeq v3 TruSeq Nano read data (paired end 301 bp, fragment length 550 bp) was downloaded from BaseSpace®, and randomly sub-sampled to ~250-fold coverage. Overlapping read pairs were merged with ABySS-mergepairs (-q 15) and resulting ca. 550 bp pseudoreads were assembled with ABySS v1.5.2 (Simpson *et al.*, 2009) (*k*=480 *l*=40 *s*=1000) yielding 67 and 61 contigs and scaffolds ≥ 500 bp, respectively. Contigs and scaffolds (Fig. S3, Table 1A and B) were scaffolded with LINKS v1.0 (*k*=15, *d*=4000, default parameters) using the *E. coli* K-12 substr. MG1655 R7 Full 2D ONT data from Quick *et al.* (2014; R7 chemistry ONI/NONI ERP007108), and results are shown in Table 1C, Fig. S4 and Table 1D, Fig. S5, in that order. SSPACE-LongRead v1.1 (*g*=200, with defaults parameters) was ran on the **B** assembly (Table 1E). ABySS scaffolds were also re-scaffolded iteratively with LINKS (*k*=15, *d*=500 to 16000, 30 iterations) using the Full 2D ONT reads (Table 1F) and, in separate experiment, all available 2D reads (Table 1G, Fig. S6) and all available R7.3 chemistry raw uncorrected reads (ERX593921;Table 1H, Fig. S7). A Baseline *S. cerevisiae* W303 Illumina MiSeq assembly (http://schatzlab.cshl.edu/data/nanocorr/) was rescaffolded with SSPACE (*g*=200), AHA (Rasko *et al.*, 2011) and LINKS (*k*=15, *d*=2-15kbp, 27 iterations) using 262,463 raw ONT reads (Fig. S8). White spruce (Genbank:ALWZ030000000, 4.2M scaffolds) was re-scaffolded with LINKS 14 times (*k*=26, *t*=200-50 *d*=5-100kbp) using draft white spruce WS77111 genotype assembly (Genbank: JZKD010000000, 4.1M sequences) (Fig. S9).

**Table 1.** QUAST analysis of a baseline *E. coli* assembly and re-scaffolded assemblies using Oxford Nanopore 2D (R7 chem.) or raw (R7.3 chem.) reads

| Stats based on sequences $\geq 500$ bp | A. ABySS Contigs | B. ABySS Scaffolds | C. LINKS k15d4000 | D. LINKS k15d4000 | E. SSPACE-LR g200 | F. LINKS 30x k15d500-16kbp | G. LINKS 30x k15d500-16kbp | H. LINKS 30x k15d500-16kbp |
| --- | --- | --- | --- | --- | --- | --- | --- | --- |
| Input Data | Illumina MiSeq | A. +MiSeq | A. +ONT Full 2D R7 | B. +ONT Full 2D R7 | B. +ONT Full 2D R7 | B. +ONT Full 2D R7 | B. +ONT All 2D R7 | B. +raw ONT R7.3 |
| Time (h:mm:ss) | - | - | 0:00:52 | 0:00:51 | 0:01:09 | 0:23:02 | 2:39:44 | 2:56:11 |
| Memory (GB) | - | - | 6.4 | 6.4 | 0.7 | 13.2 | 85.4 | 104.8 |
| Total sequences | 67 | 61 | 49 | 48 | 43 | 27 | 16 | 27 |
| Largest (bp) | 358,719 | 406,793 | 633,204 | 633,147 | 628,411 | 1,057,556 | 1,286,148 | 1,286,419 |
| NG50 length (bp) | 179,720 | 204,529 | 270,992 | 270,992 | 226,696 | 633,147 | 1,197,321 | 645,797 |
| Misassemblies | 5 | 5 | 5 | 5 | 8 | 11 | 20 | 9 |
| Genes + parts | 4,442 + 63 | 4,443 + 62 | 4,443 + 62 | 4,443 + 62 | 4,448 + 57 | 4,443 + 62 | 4,440 + 62 | 4,443 + 62 |
| Max alignment (bp) | 358,223 | 405,659 | 486,572 | 486,572 | 405,659 | 760,934 | 760,934 | 759,131 |
| NGA50 (bp) | 177,531 | 179,569 | 228,879 | 228,879 | 226,324 | 299,206 | 299,206 | 486,527 |

### 2.2 LINKS Algorithm

#### Process

ONT reads are supplied as input (*-s* option, fasta format) and *k-mer* pairs are extracted using user-defined *k-mer* length (*-k*) and distance between the 5’-end of each pairs (*-d*) over a sliding window (*-t*). Unique *k-mer* pairs at set distance are hashed. Fasta sequences to scaffold are supplied as input (*-f*), and are shredded to *k-mers* on both strands, tracking the contig or scaffold of origin, *k-mer* positions and frequencies of observation.

#### Algorithm

LINKS has two main stages: contig pairing, and scaffold layout. Cycling through *k-mer* pairs, *k-mers* that are uniquely placed on contigs are identified. Putative contig pairs are formed if *k-mer* pairs are on different contigs. Contig pairs are only considered if the calculated distances between them satisfy the mean distance provided (*-d*), while allowing for a deviation (*-e*). Contig pairs having a valid gap or overlap are allowed to proceed to the scaffolding stage. Contigs in pairs may be ambiguous: a given contig may link to multiple contigs. To mitigate, the number of spanning *k-mer* pairs (links) between any given contig pair is recorded, along with a mean distance estimate. Once pairing between contigs is complete, the scaffolds are built using contigs in turn until all have been incorporated into a scaffold. Scaffolding is controlled by merging sequences only when a minimum number of links (*-l*) join two contig pairs, and when links are dominant compared to that of another possible pairing (*-a*). The predecessor of LINKS is the unpublished scaffolding engine in the widely-used SSAKE assembler (Warren *et al.*, 2007), and foundation of the SSPACELongRead scaffolder (Boetzer and Pirovano, 2014).

#### Output

A summary of the scaffold layout is provided (.scaffold) as a text file, and captures the linking information of successful scaffolds. A fasta file (.scaffold.fa) is generated using that information, placing N-pads to represent the estimated lengths of gaps, and a single “n” in cases of overlaps between contigs. A log summary of *k-mer* pairing in the assembly is provided (.log) along with a text file describing possible issues in pairing (.pairing_issues) and pairing distribution (.pairing_distribution.csv).

## 3 RESULTS

We used a publicly available ONT data for scaffolding *E. coli* contigs and scaffolds derived from high-depth Illumina MiSeq 300 bp paired end reads and assembled with ABySS. These assemblies were already very contiguous (scaffold NG50=204 kbp) and of good quality, as assessed by QUAST (Gurevich *et al.*, 2013, Table 1). First, we scaffolded ABySS contigs with only the ONT Full 2D data (*k*=15, *d*=4000), as a benchmark of the method, yielding an improved assembly that rivaled the ABySS scaffolder on Illumina data (Table 1B) in assembly quality (Table 1C). The runs were fast (<1 min), and required moderate resources (~6 GB RAM). Next, we re-scaffolded *E. coli* ABySS scaffolds (**B.**) using the same parameters. Running SSPACE-LongRead (**E.**) gave similar results, but using less memory by a factor 10. Further merge opportunities are found by comprehensively extracting paired *k-mers* at various distance intervals. Accordingly, we ran LINKS iteratively 30 times, each instance increasing the distance between *k-mer* pairs from 500 to 16,000 bp and gradually improving the long-range contiguity of the assembly (Table 1F and Fig. S5). Despite the relatively longer runtime (23 min.), the final assembly was in less than half the original number of sequences, and the scaffold NG50 length exceeded 600 kbp. With the same run parameters, using all available 2D ONT data, LINKS required 85 GB RAM and 2h39m to complete (Table 1G), and yielded 12 scaffolds >1 kbp that are co-linear with the *E. coli* genome (Fig. S6). Using raw R7.3 ONT reads for scaffolding gave the best compromise between errors and contiguity (Table 1H). We tested LINKS on the larger *S. cerevisiae* ONT dataset (Goodwin *et al.*, 2015) and obtained an assembly that compares with the Celera Assembly of Illumina-corrected ONT reads (Nanocorr) in contiguity, but with less than half the errors (Fig. S8). High RAM usage with LINKS can be mitigated by increasing the sliding window step (*-t*), which decreases the *k-mer* pair space. Doing so, re-scaffolding the colossal white spruce genome draft assembly (Birol *et al.*, 2013) 14 times using the draft assembly of another white spruce genotype was possible with ~ 132 GB RAM, producing a conifer assembly whose NG50 contiguity is in excess of 110 kbp (Fig. S9).

To our knowledge, LINKS is the first publicly available scaffolder designed specifically for nanopore long reads and with a general framework that could apply to scaffolding very large genomes, such as that of white spruce (20 Gbp) using another assembly draft or reference in lieu of long reads. This study highlights the present utility of ONT reads for genome scaffolding in spite of their current limitations, which are expected to diminish as the nanopore sequencing technology advances.

## ACKNOWLEDGEMENTS

This work is partly funded by Genome Canada, British Columbia Cancer Foundation, and Genome British Columbia. Research reported in this publication was also partly supported by the National Human Genome Research Institute of the National Institutes of Health under Award Number R01HG007182. The content is solely the responsibility of the authors and does not necessarily represent the official views of the National Institutes of Health or other funding organizations.

## SUPPLEMENTARY MATERIAL

**Supplementary Figure S1.**
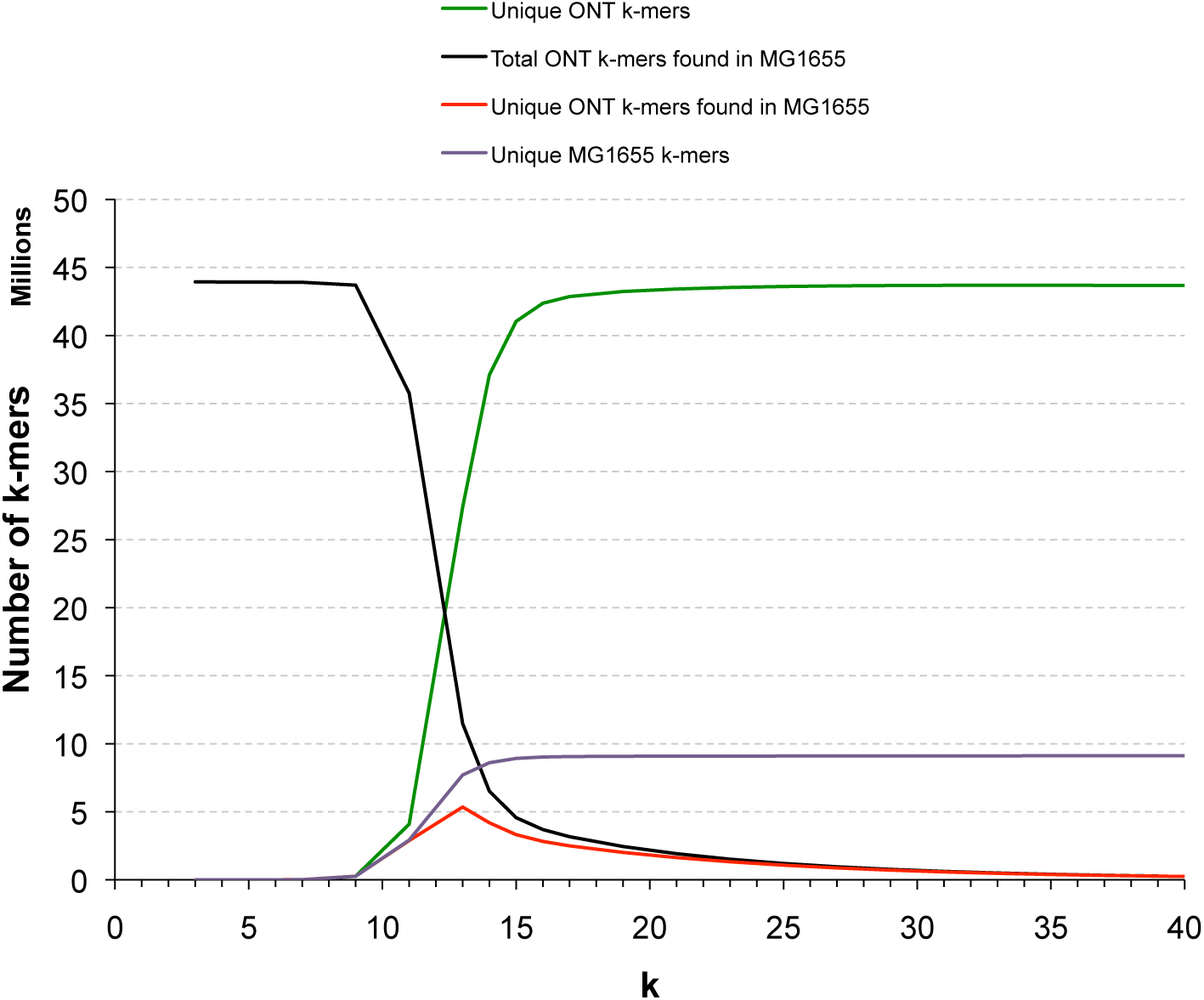
Full 2D ONT long read *k-mer* uniqueness in the *E. coli* K-12 genome reference. *k-mers* were extracted from both the Full 2D ONT data (Quick *et al.*, 2014) and the *E. coli* K-12 substr. MG1655 (accession U00096.2) reference genome sequence. A Bloom filter (Bloom, 1970) was built from the latter and *k-mers* extracted from the former files used to query the filter for matching sequences. *k*=15 gives the best compromise of specificity, yield and uniqueness with the data set at hand.

**Supplementary Figure S2.**
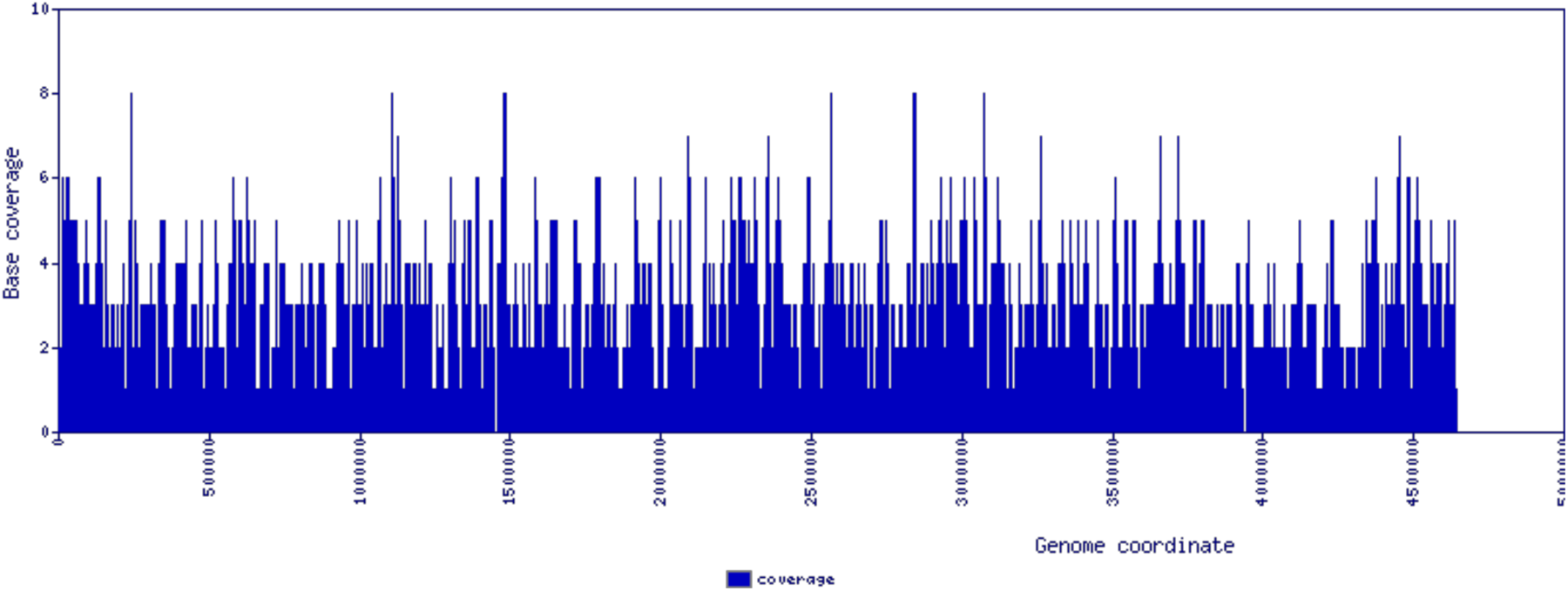
*E. coli* K-12 substr. MG1655 genome coverage analysis by Full 2D (R7 chemistry) Oxford Nanopore long reads. High-quality 2D nanopore reads (Quick et al. 2014) were aligned with blastn (Altshul et al. 1990) onto the *E. coli* K-12 substr. MG1655 reference (U00096.2), plotting only reads with sequence identity over 50% (1,713 high quality sequences out of 3,471). We identified 184 regions 1 bp and longer with no read coverage. Overall 90.3% of the 4,639,675 bp MG1655 genome was covered by at least one nanopore read.

**Supplementary Figure S3.**
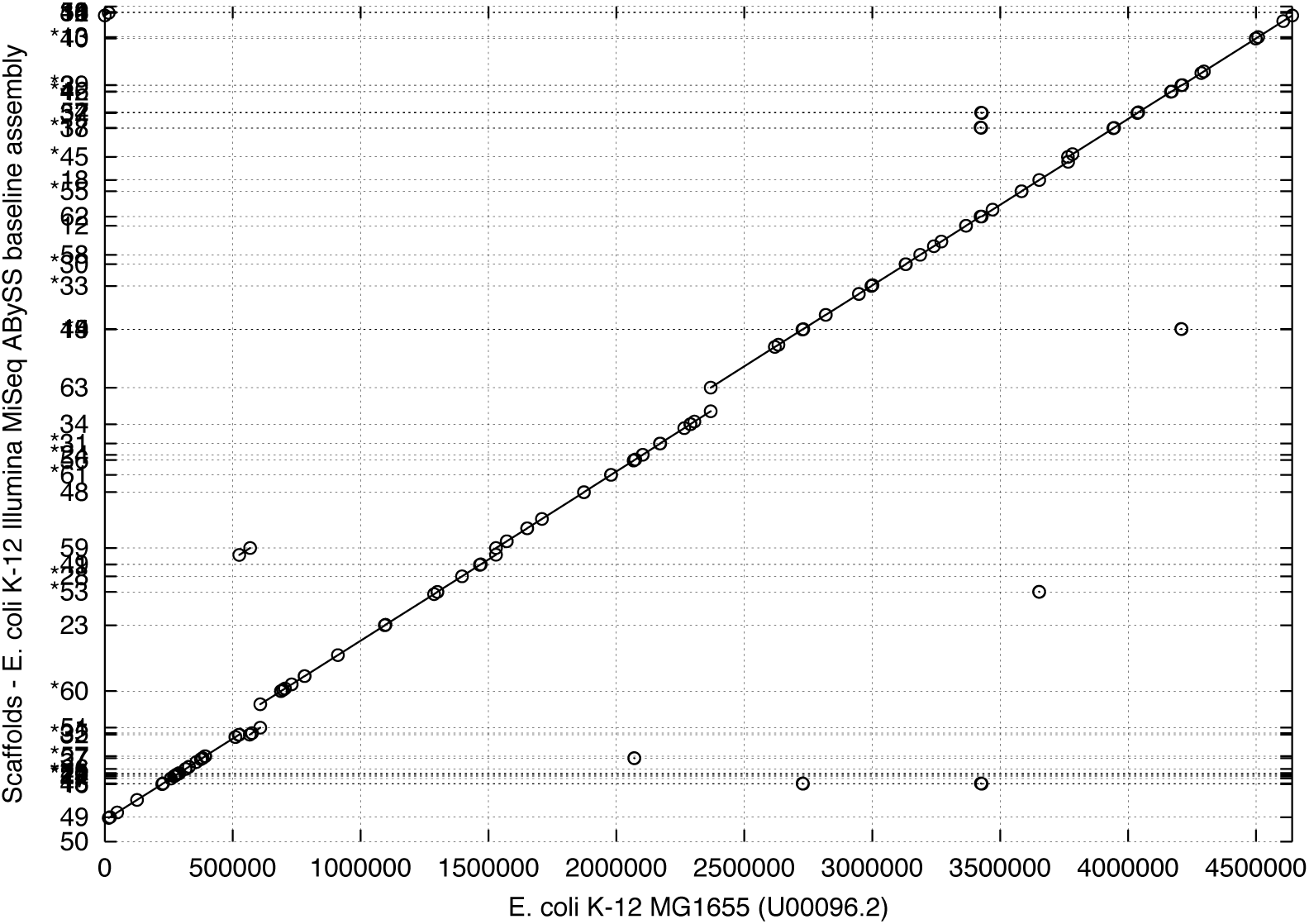
*E. coli* K-12 assembly and genome co-linearity. A baseline ABySS assembly (Table 1B in main text) of the *E. coli* K-12 MG1655 genome yields a draft genome that despite being fragmented is co-linear with the reference. Sequence comparison was performed with MUMmer v3.23 tools, using nucmer for nucleotide sequence alignments and mummerplot for plotting (Kurtz *et al.*, 2004).

**Supplementary Figure S4.**
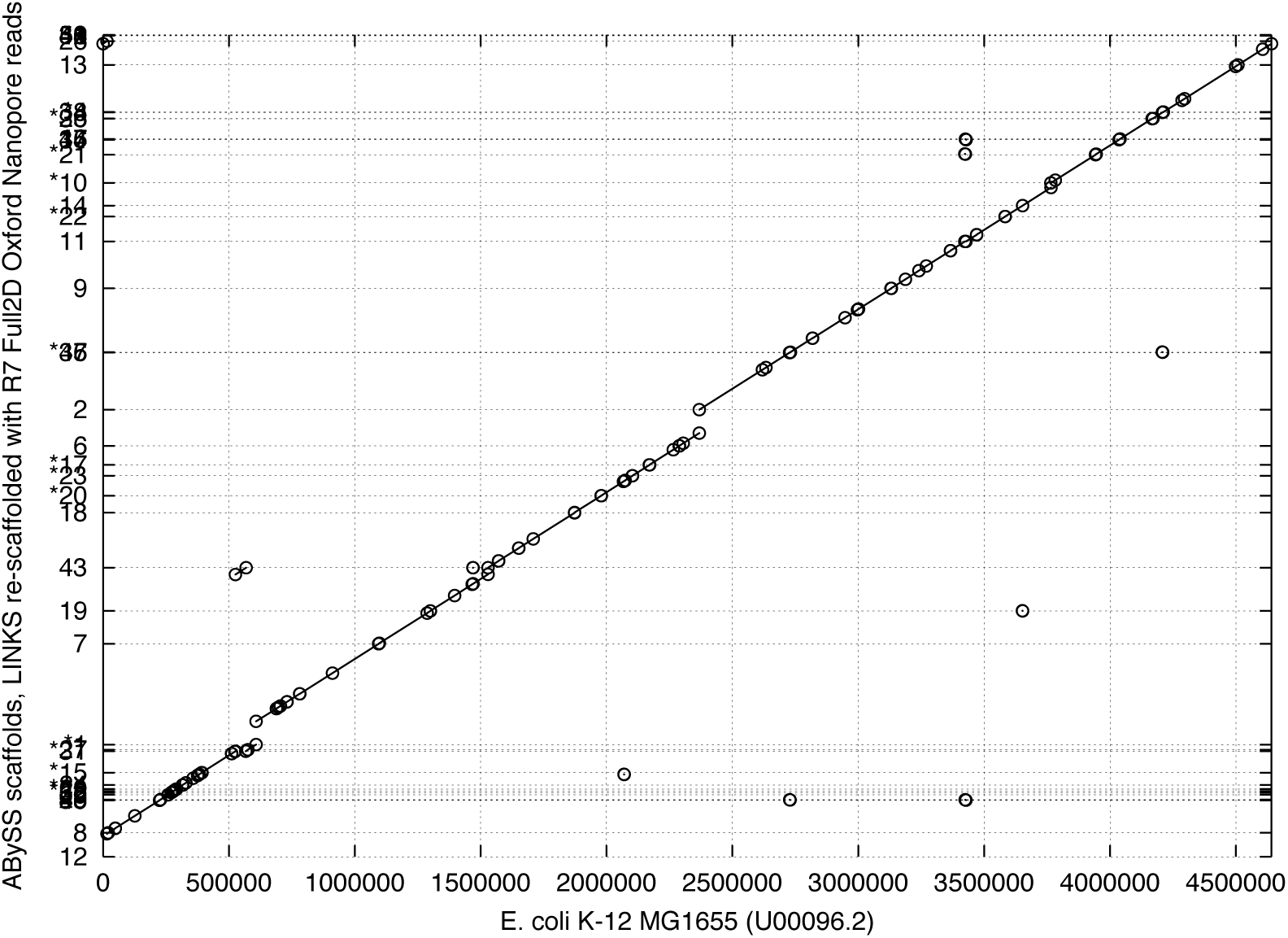
Full 2D ONT - LINKS scaffolds co-linearity with the MG1655 genome, single *k-mer* pair LINKS run. A single LINKS scaffolding round (k=15 bp, d=4000 bp) was performed on ABySS assembly sequence scaffolds (shown in Fig. S3B), bringing the number of scaffolds from 61 to 48 (Table 1D in manuscript) and harboring sequences in the correct order and orientation.

**Supplementary Figure S5.**
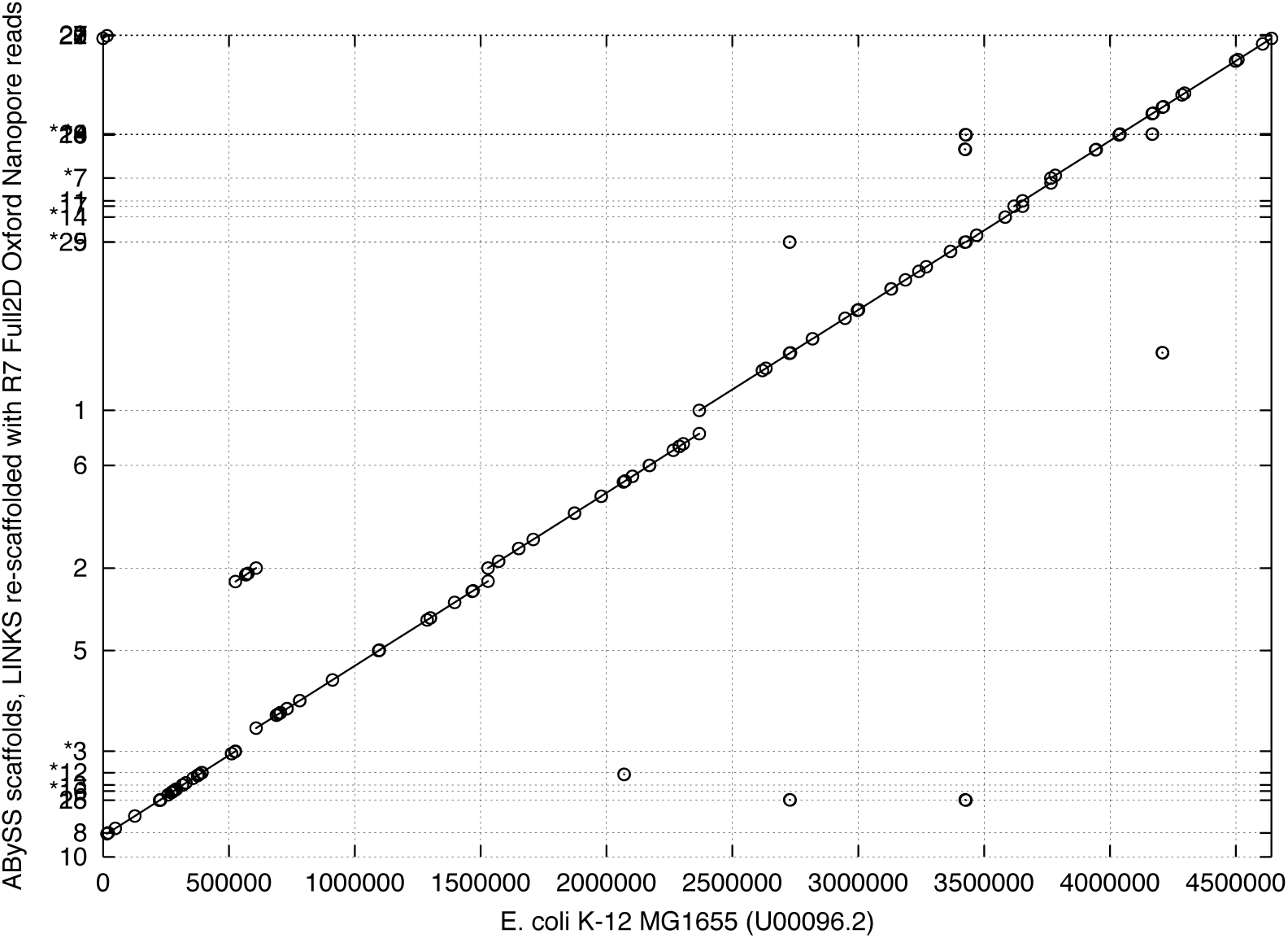
Full 2D ONT-LINKS scaffolds co-linearity with the reference *E. coli* K-12 genome (multiple *k-mer* pair runs). Iterative LINKS scaffolding rounds (*k*=15, *d*=500 to 16000 bp, 30 iterations) were performed on ABySS assembly sequence scaffolds (Table 1F in manuscript), bringing the number of scaffolds further down to 27 from 61, with its underlying sequences in the exact configuration compared to the reference.

**Supplementary Figure S6.**
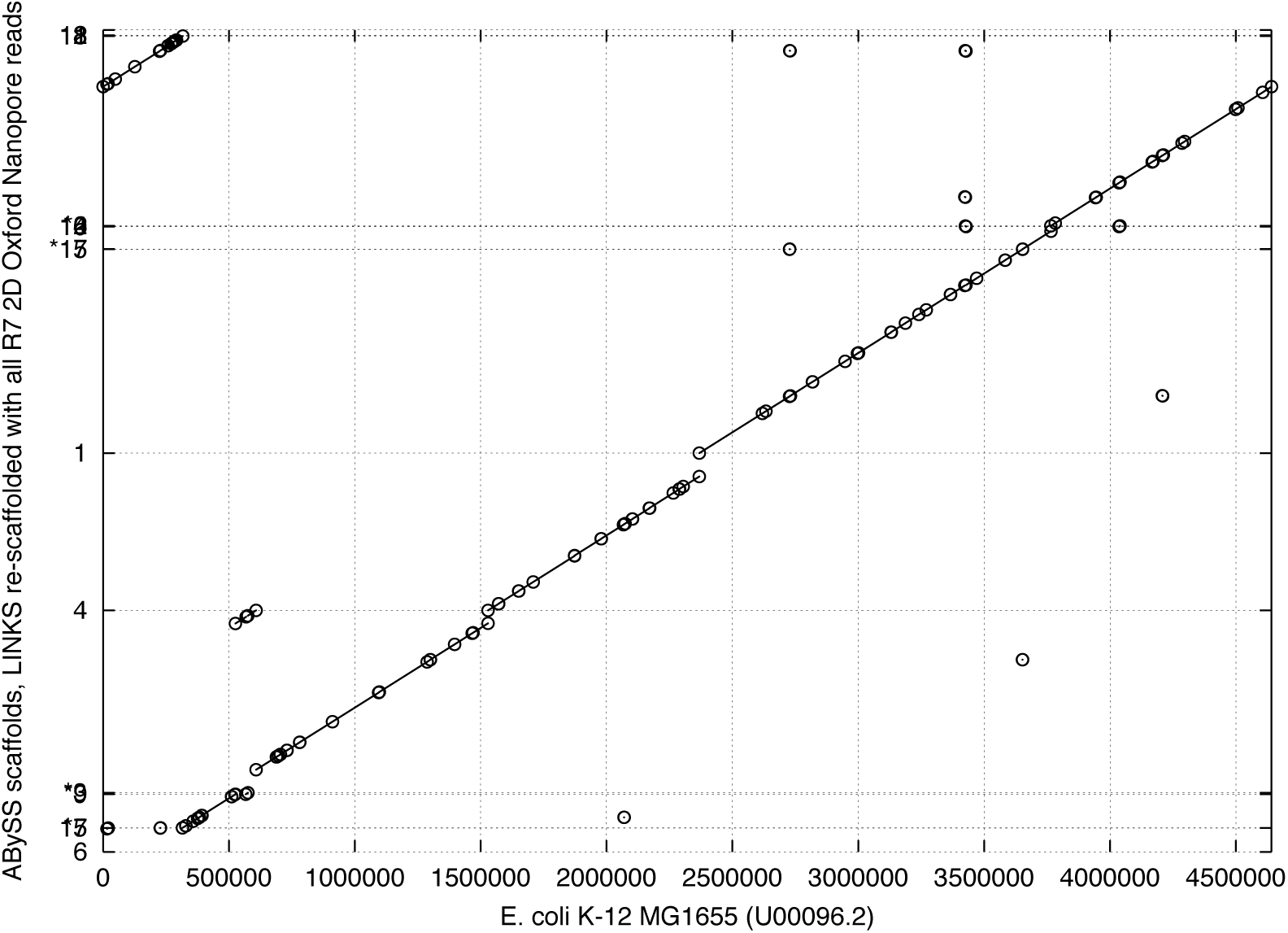
LINKS scaffolds using all available R7 2D ONT reads compared to the reference *E. coli* K-12 genome (forty five *k-mer* pair interval runs). Iterative LINKS scaffolding rounds (*k*=15, *d*=500 to 16000 bp, 30 iterations) were performed on ABySS assembly sequence scaffolds (Table 1G in manuscript), bringing the number of scaffolds further down to 16 from 61. MUMmer co-linear analysis indicates that six large scaffolds comprise *E. coli* K-12 MG1655 re-scaffolded sequences in the correct order and orientation.

**Supplementary Figure S7.**
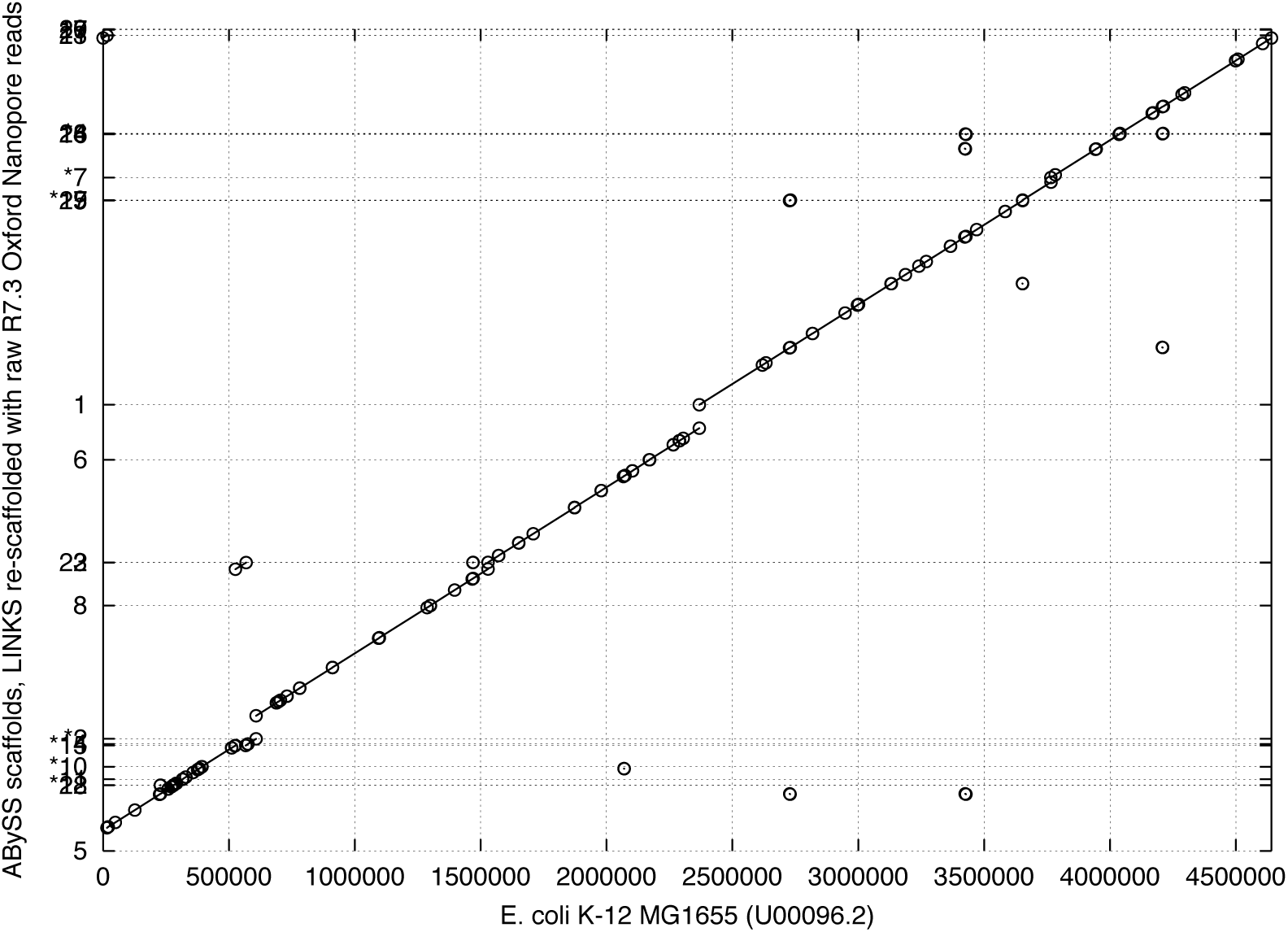
LINKS scaffolds using all raw, uncorrected R7.3 ONT reads compared to the reference *E. coli* K-12 genome (forty five k-mer pair interval runs). Iterative LINKS scaffolding rounds (k=15, d=500 to 16000 bp, 30 iterations) were performed on the baseline ABySS assembly sequence scaffolds (Table 1H in manuscript), bringing the number of scaffolds down to 27 from 61. QUAST analysis reveals that re-scaffolding with the raw v7.3 ONT data produces an assembly with the best compromise, with fewer errors and highest overall contiguity.

**Supplementary Figure S8.**
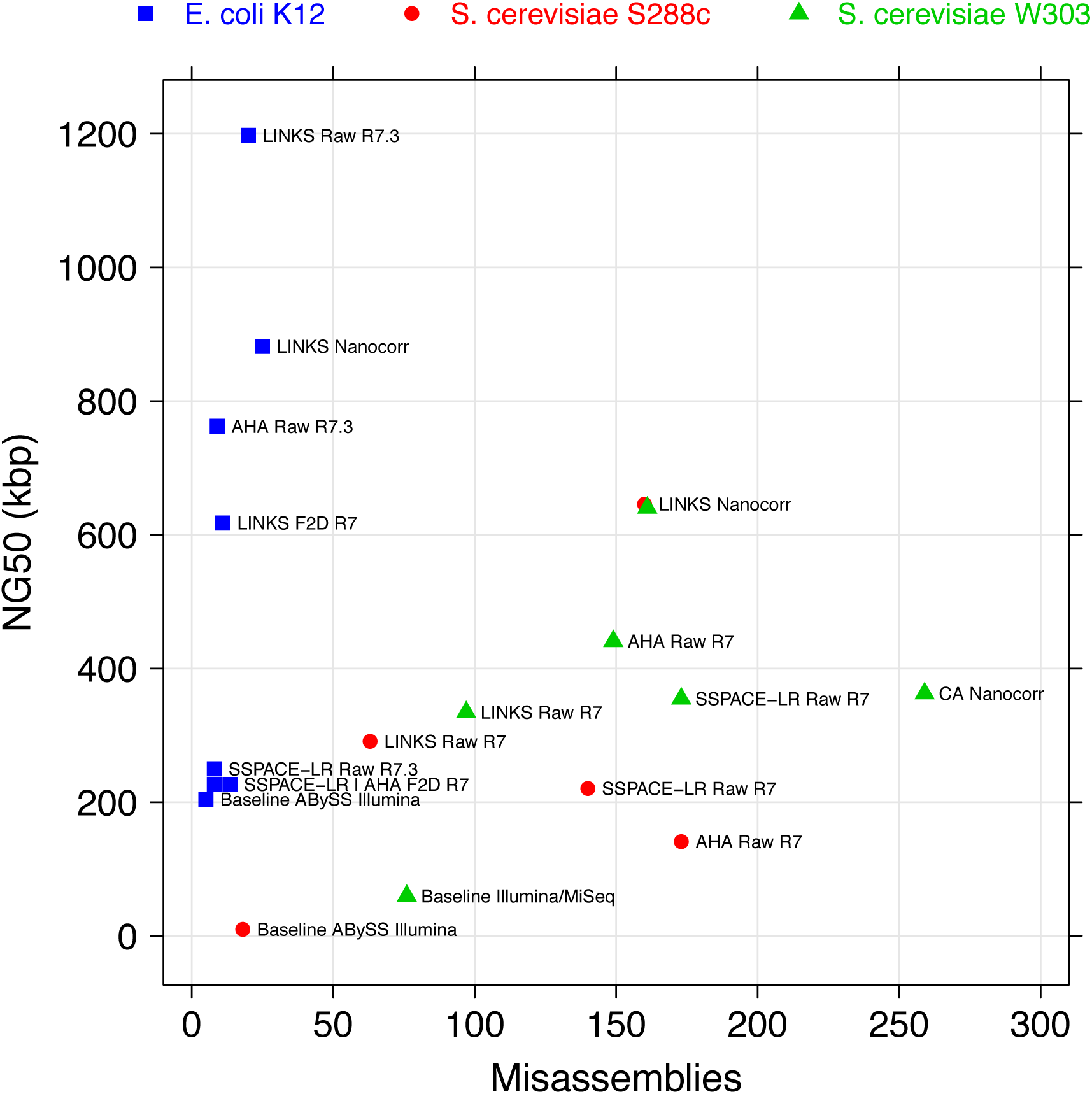
Scaffolding high-quality short read assemblies with Oxford Nanopore Technologies long reads. Publicly available ONT long reads for *E. coli* K12 MG1655 and *S. cerevisiae* W303 were recently made available (Quick et al. 2014; Goodwin et al 2015). We have used these data to re-scaffold baseline ABySS scaffold assemblies of Illumina-only data (Simpson *et al.*, 2008; Base space: Illumina MiSeq v3 TruSeq Nano read data and SRA:ERR156523) using LINKS, AHA (Rasko *et al.*, 2011) and SSPACE-LR (Boetzer and Pirovano 2014) (blue squares and red circles). Additionally, we have re-scaffolded a baseline Illumina MiSeq assembly of *S. cerevisiae* W303 (http://schatzlab.cshl.edu/data/nanocorr/) with the same tools and compared it to a Celera Assembler assembly of Illumina-corrected ONT reads (green triangles). The *E.coli* Nanocorr CA assembly was excluded from the graph for clarity; it produced a single scaffold with only 5 errors. ABYSS Illumina, Illumina-only scaffold assemblies; SSPACE-LR F2D R7, SSPACE-LR scaffolding of the base assembly using full 2D ONT reads R7 chemistry; LINKS F2D R7, LINKS 30x iterative scaffolding using full 2D ONT reads R7 chemistry; LINKS Raw R7.3, LINKS 30x iterative scaffolding using raw ONT reads R7.3 chemistry; LINKS Raw R7, LINKS iterative scaffolding using raw W303 ONT reads to rescaffold a S288c ABySS assembly (31 iterations, purple circle) or a W303 Illumina/MiSeq assembly (27 iterations, green triangle); SSPACE-LR Raw R7, SSPACE-LR scaffolding of the base assembly (S288c ABySS illumina purple circle and W303 Illumina MiSeq green triangle) using raw W303 ONT reads); CA Nanocorr, Celera Assembler assembly of Illumina-corrected W303 ONT reads; AHA, Pacific Biosciences’ A Hybrid Assembler; LINKS Nanocorr, 27x *(E.coli* K12), 28x *(S. cerevisiae* W303) or 39x *(S. cerevisiae* S288c) LINKS scaffolding iterations of baseline assemblies using respective Nanocorr corrected reads (http://schatzlab.cshl.edu/data/nanocorr/).

**Supplementary Figure S9.**
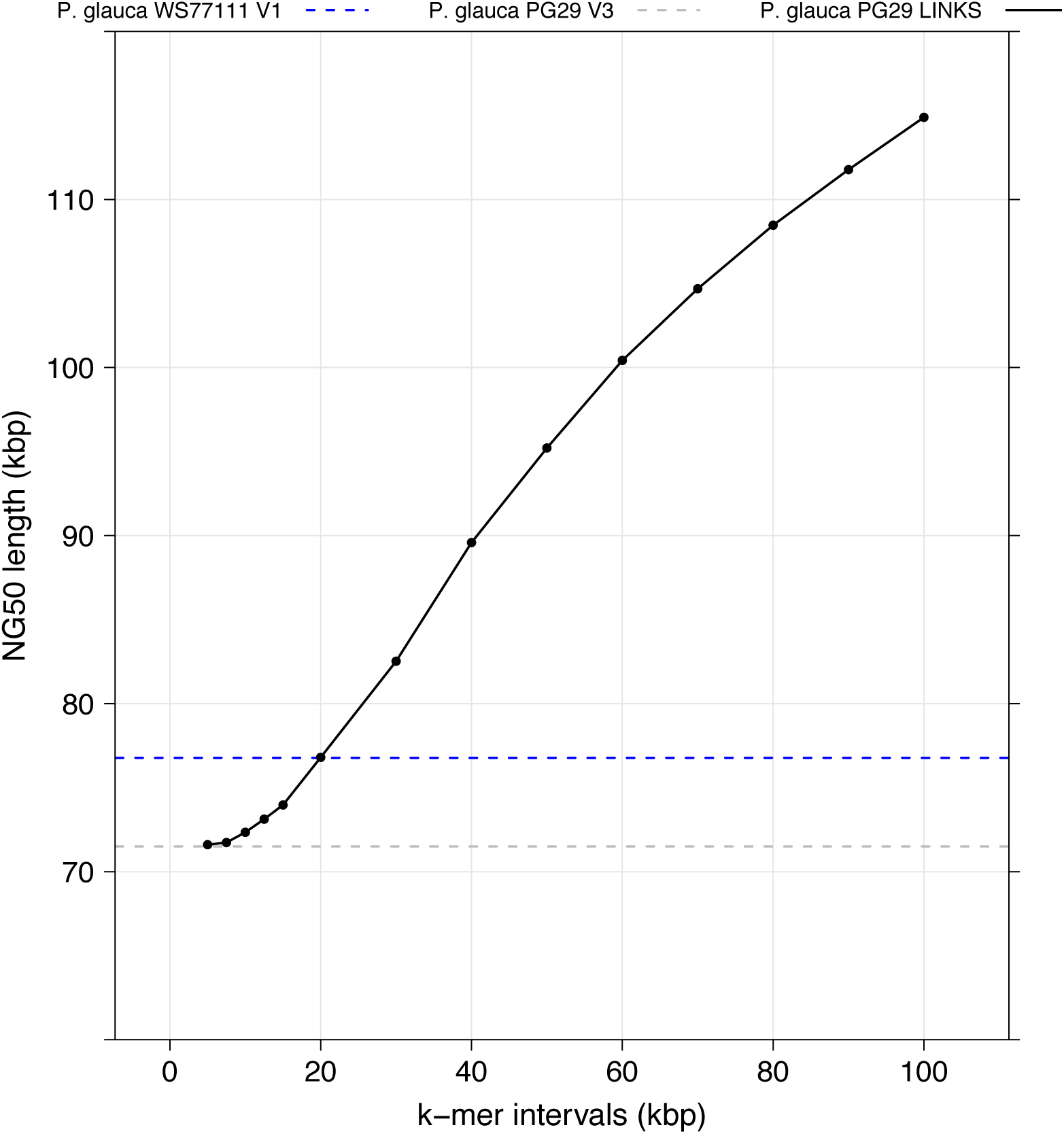
LINKS re-scaffolding of the white spruce (*P. glauca,* PG29 cultivar) genome with *k-mer* pairs derived from the white spruce WS77111 genotype draft assembly. Iterative LINKS scaffolding rounds (fourteen iterations, *k*=26*, t*=200 to 50, *l*=5, *a*=0.3, *d*=5 kbp to 100kbp, interval shown on *x* axis, solid black line) were performed on the PG29 V3 ABySS assembly sequence scaffolds (Genbank: ALWZ030000000, 4.2M scaffolds ≥ 500bp, dotted grey line, Birol *et al.*, 2013) using sequence data from the WS77111 V1 draft assembly (Genbank:JZKD010000000, 4.3M scaffolds ≥ 500bp, dotted blue line), increasing the PG29 assembly contiguity 1.5-fold to reach an NG50 length (Earl *et al.*, 2011) of 114,888 bp (4.1M scaffolds ≥ 500bp).

